## Supplementary figures and images for "Numb provides a fail-safe mechanism for intestinal stem cell self-renewal in adult *Drosophila* midgut"

### Figure-5-figure supplement1

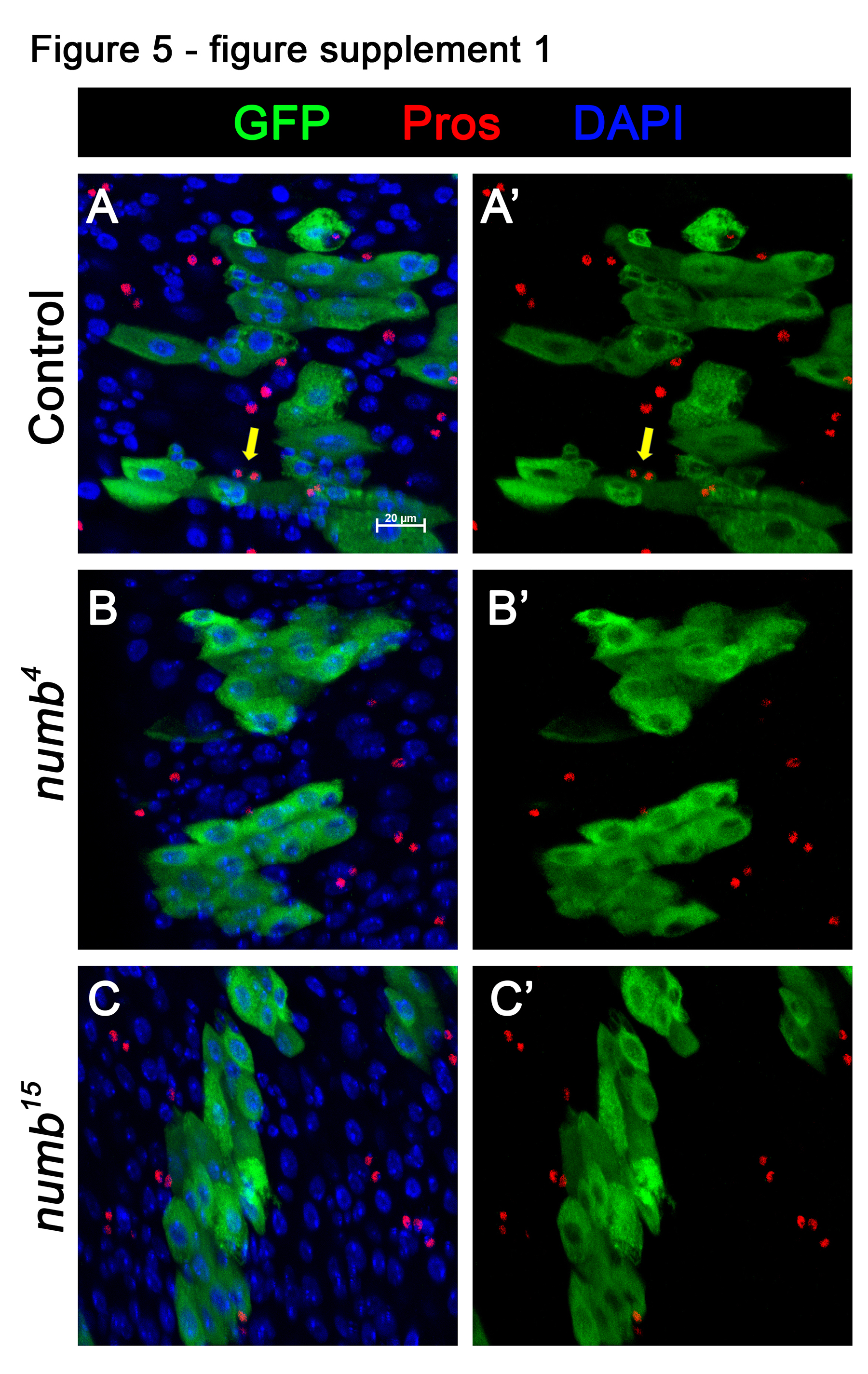
